# JGI Plant Gene Atlas: An updateable transcriptome resource to improve structural annotations and functional gene descriptions across the plant kingdom

**DOI:** 10.1101/2022.09.30.510380

**Authors:** Avinash Sreedasyam, Christopher Plott, Md Shakhawat Hossain, John T. Lovell, Jane Grimwood, Jerry W. Jenkins, Christopher Daum, Kerrie Barry, Joseph Carlson, Shengqiang Shu, Jeremy Phillips, Mojgan Amirebrahimi, Matthew Zane, Mei Wang, David Goodstein, Fabian B. Haas, Manuel Hiss, Pierre-François Perroud, Sara S. Jawdy, Rongbin Hu, Jenifer Johnson, Janette Kropat, Sean D. Gallaher, Anna Lipzen, Ryan Tillman, Eugene V. Shakirov, Xiaoyu Weng, Ivone Torres-Jerez, Brock Weers, Daniel Conde, Marilia R. Pappas, Lifeng Liu, Andrew Muchlinski, Hui Jiang, Christine Shyu, Pu Huang, Jose Sebastian, Carol Laiben, Alyssa Medlin, Sankalpi Carey, Alyssa A. Carrell, Mariano Perales, Kankshita Swaminathan, Isabel Allona, Dario Grattapaglia, Elizabeth A. Cooper, Dorothea Tholl, John P. Vogel, David J Weston, Xiaohan Yang, Thomas P. Brutnell, Elizabeth A. Kellogg, Ivan Baxter, Michael Udvardi, Yuhong Tang, Todd C. Mockler, Thomas E. Juenger, John Mullet, Stefan A. Rensing, Gerald A. Tuskan, Sabeeha S. Merchant, Gary Stacey, Jeremy Schmutz

## Abstract

Gene functional descriptions, which are typically derived from sequence similarity to experimentally validated genes in a handful of model species, offer a crucial line of evidence when searching for candidate genes that underlie trait variation. Plant responses to environmental cues, including gene expression regulatory variation, represent important resources for understanding gene function and crucial targets for plant improvement through gene editing and other biotechnologies. However, even after years of effort and numerous large-scale functional characterization studies, biological roles of large proportions of protein coding genes across the plant phylogeny are poorly annotated. Here we describe the Joint Genome Institute (JGI) Plant Gene Atlas, a public and updateable data resource consisting of transcript abundance assays from 2,090 samples derived from 604 tissues or conditions across 18 diverse species. We integrated across these diverse conditions and genotypes by analyzing expression profiles, building gene clusters that exhibited tissue/condition specific expression, and testing for transcriptional modulation in response to environmental queues. For example, we discovered extensive phylogenetically constrained and condition-specific expression profiles across many gene families and genes without any functional annotation. Such conserved expression patterns and other tightly co-expressed gene clusters let us assign expression derived functional descriptions to 64,620 genes with otherwise unknown functions. The ever-expanding Gene Atlas resource is available at JGI Plant Gene Atlas (https://plantgeneatlas.jgi.doe.gov) and Phytozome (https://phytozome-next.jgi.doe.gov), providing bulk access to data and user-specified queries of gene sets. Combined, these web interfaces let users access differentially expressed genes, track orthologs across the Gene Atlas plants, graphically represent co-expressed genes, and visualize gene ontology and pathway enrichments.

## INTRODUCTION

The flowering plant, *Arabidopsis thaliana*, has served as a model for functional genomics over the past two decades. While the goal of functionally characterizing each *A. thaliana* gene by the year 2010 (Koornneef and Meinke 2010) has yet to be fully realized, many large-scale studies, such as gene knock-out collections for reverse genetics, have tested the phenotypic effects nearly half of *A. thaliana* protein-coding genes (Berardini et al. 2015). These experimentally validated loci, and a massive set of predicted and curated gene functions form the foundation for gene characterization across 400M years of plant evolution.

Despite the potential for homology-based functional annotations across plants, putative gene functions in non-model plants are sparse, often containing a majority of genes with no functional descriptions. These knowledge gaps are undoubtedly due to the phylogenetic and functional scale of plant diversity. At one extreme, DNA or protein sequences may have diverged so that no genes have obvious *A. thaliana* homologs. However, even with homology, assigning gene function to distantly related plants assumes function is evolutionarily conserved. This assumption is clearly violated in many situations: flowering plants have evolved diverse adaptive traits, specialized organs/tissues, and environmental responses, all of which are poorly captured by a single model organism. Further, gene neofunctionalization, subfunctionalization and gene cooption may invalidate direct superimposition of gene annotation from one species to another (C. Li et al. 2012; Nicotra et al. 2010; Raissig et al. 2017). The addition of other model species, including *Brachypodium distachyon, Oryza sativa*, and *Physcomitrium patens*, has helped fill gaps in homology-based functional annotations. However, 16.1-56.9% (*M*= 27.8; *SD* = 10.06) of protein coding genes across the plant phylogeny remain poorly characterized (**Supplemental Fig. 1**) (Gollery et al. 2006, 2007; Rhee and Mutwil 2014).

Incomplete gene functional annotations are not only due to an overreliance on few genetic model organisms, but also an inability to link experimental evidence across species. However, centralized functional databases containing information generated from new experiments such as ongoing large-scale transcriptome projects and genome-wide association studies could accelerate gene function discovery. Even with a central repository, interpretation and integration across diverse studies is difficult because experimental and analytical protocols are rarely standardized. For example, different sample collection, RNA isolation, library construction protocols, and sequencing platforms can result in significant variation in sequence coverage and estimates of gene expression (Levin et al. 2010; Ross et al. 2013; Sudmant, Alexis, and Burge 2015; Yu et al. 2014). This among-experiment variation reduces the accuracy and precision of comparisons across species and studies, which directly limits putative gene function inference from transcript abundance profiles.

Here, we present an updateable large-scale dataset and a suite of experimental protocols to facilitate functional gene prediction across the diversity of plants. Crucially, we have developed experimental conditions, tissue types, and analytical protocols that permit comprehensive analysis of gene expression across plants. We applied these conditions and collected 2,090 tissue samples from 18 plant species spanning single-celled algae, bryophytes, and flowering plants. This integrated dataset (1) forms a foundation to improve gene functional annotations, (2) facilitates cross-species comparative transcriptomics within controlled environmental and laboratory conditions, and (3) permits high-powered tests of gene regulatory evolution across phylogenetically diverse plant genomes. To demonstrate this functionality, we cataloged the expression profiles of annotated genes, and built co-expressed clusters of genes that exhibited tissue/condition specific expression patterns including responses to changes in nitrogen (N) regimes, abiotic stressors, and developmental stages. We systematically assigned expression derived functional descriptions to an average of 40.6% (*SD* = 12.6) of annotated genes in the assessed genomes, 9.5% of which previously had no known function. This substantial transcriptomic resource is available to the research community at JGI Plant Gene Atlas (https://plantgeneatlas.jgi.doe.gov) and through Phytozome, the JGI Plant Portal, at https://phytozome-next.jgi.doe.gov (Goodstein et al. 2012).

## SCOPE OF DATA GENERATED

We developed the JGI Gene Atlas from 15.4 trillion sequenced RNA bases (Tb) and 2,090 RNA-seq samples across 9 JGI plant flagship genomes and 9 other reference plants (**Table 1**). For each of the sequenced plants, we collected tissue samples representing appropriate developmental stages, growth conditions, tissues, and abiotic stresses (**Fig. 1**). To reduce residual environmental variance, we followed standard growth conditions including light quality, quantity and duration, temperature, water, growth media, and nutrients. Experimental treatments were applied using standardized methods across all species (see Methods).

**Table 1.**
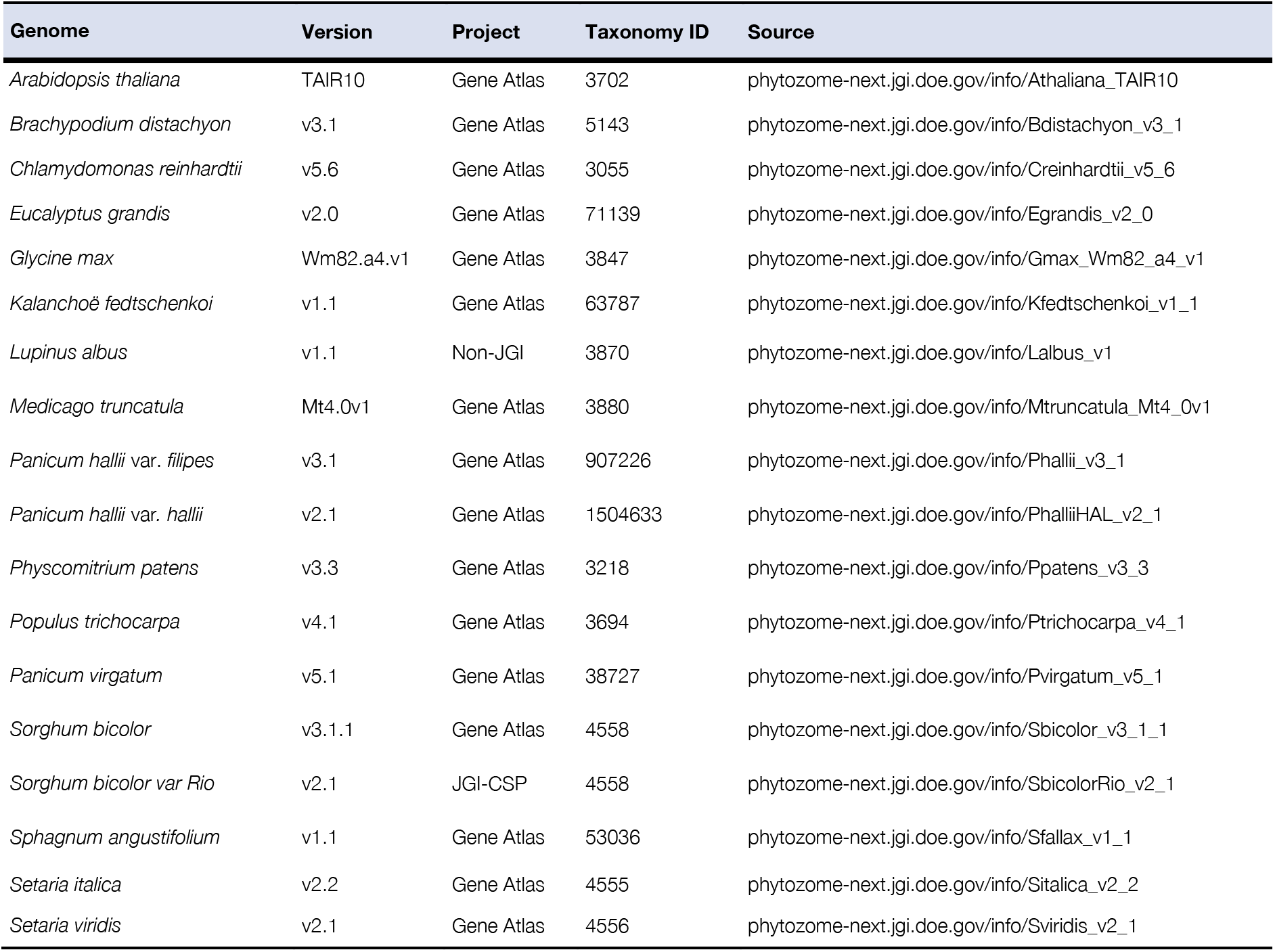
JGI Plant Gene Atlas species. Genome annotation versions of 18 diverse plants included in the current release.

**Figure 1.**
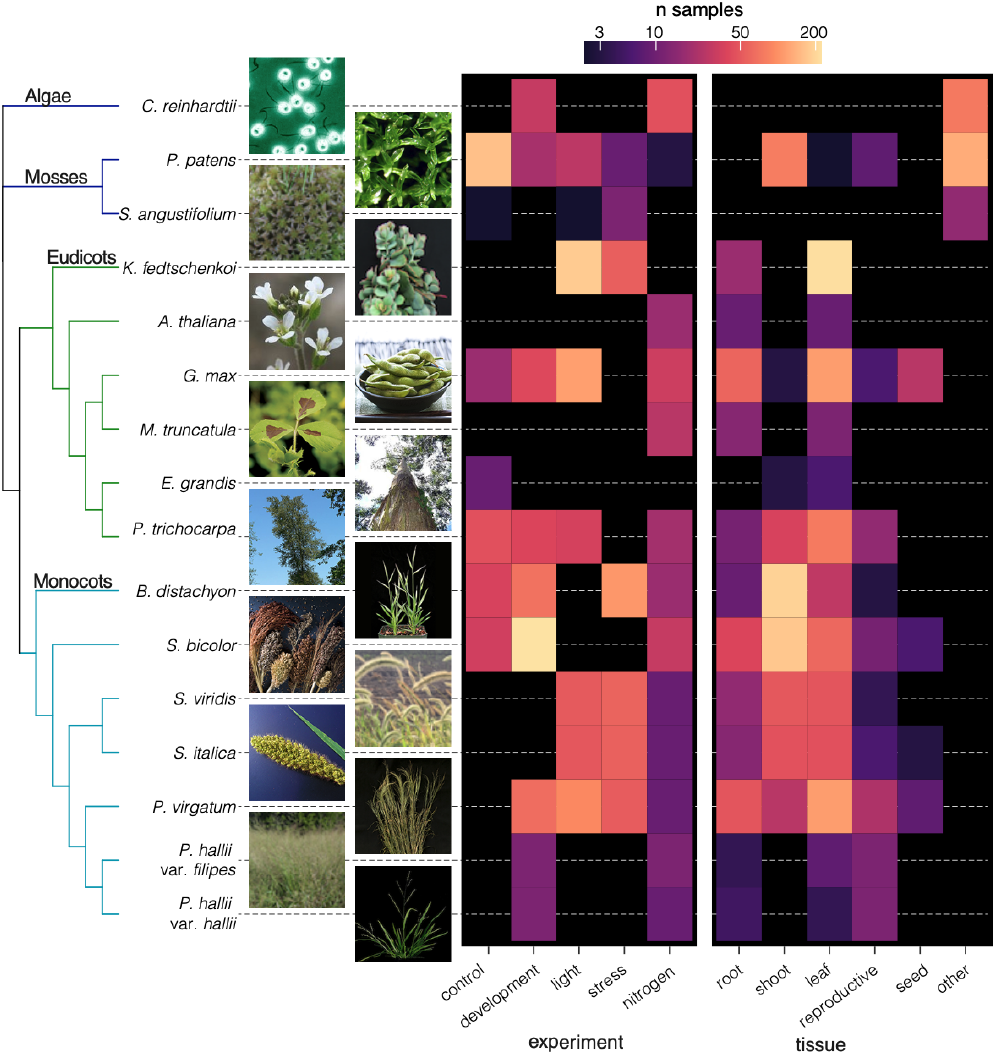
The phylogenetic context and scope of Gene Atlas RNA-seq samples. The 16 genomes are ordered by their phylogenetic position, visualized on the left as a cladogram without branch lengths that was constructed from 10 single-copy orthologs. Tips are labeled with genome names and thumbnail photos. Photo credit given on Phytozome.

We sought to limit among-experiment measurement and environmental variation by using identical molecular methods to extract (RNA integrity number, RIN ≥ 5 and at least 1 μg of total RNA) and sequence (Illumina stranded, paired-end 2×150 RNA-seq libraries) high-quality RNA. All samples were quality tested and sequenced at JGI. The resulting transcript abundance assays were highly correlated across biological replicates within conditions, tissues, and genotypes (**Supplemental Data 1**), which provides evidence that our gene expression measurements are highly accurate and robust.

We also demonstrated that the JGI Gene Atlas is updateable, with a new reference genome version and even with sequence data derived from other experiments and sequencing facilities. To accomplish this, we included *S. bicolor* ‘Rio’ (sweet sorghum, n_samples_ = 94) (Cooper et al. 2019) from JGI’s Community Science Program project and *Lupinus albus* (white lupin) cluster root tissue (n_samples_ = 72) (Hufnagel et al. 2020) from a non-JGI project. A comprehensive list of all samples available so far is in **Supplemental Data 2** and https://plantgeneatlas.jgi.doe.gov. Our custom pipeline to analyze expression levels of protein-coding genes is outlined in **Supplemental Fig. 2**.

## OVERVIEW OF THE TRANSCRIPTOMIC LANDSCAPE OF GENE ATLAS PLANTS

### Developing a baseline of evolutionarily conserved gene expression

Across all 18 species, 47-87% (mean = 73%) of annotated genes were transcriptionally active (FPKM > 1). To test for conserved and divergent expression levels across the 18 species, we applied the traditional method of comparing single-copy orthologs across species. While powerful, restricting tests to orthologs based on gene sequences can be problematic across evolutionarily diverged lineages. For example, given the phylogenetic distance and nested whole-genome duplications among our sampled species, we were only able to find 2,066 one-to-one orthologous protein-coding genes (**Supplemental Data 3**) across just eight of the vascular plant genomes. Furthermore, such single-copy orthologs have evolutionarily conserved sequences and likely gene functions, permitting better homology-based functional descriptions (89.01% with good functional descriptions) than genome-wide averages (83.8%, Fisher’s exact test odds = 1.607, *P* = 5.495e-12). Nonetheless, we observed 227 (10.98%) genes with 1:1 orthologs and consistent expression among species, but weak functional descriptions (**Supplemental Data 4**). Given the expected paucity of multi-genome single-copy orthologs, we also addressed the challenge of finding genes with similar expression across species by analyzing pairwise single-copy orthologs to a single reference genome, *A. thaliana*. Overall, we identified 6,018 unique *Arabidopsis* orthologs that showed conserved expression patterns across multiple species. Surprisingly, these genes include 660 (11%) with little to no known functional description, making these genes rational targets for functional characterization studies (**Supplemental Data 5**). Identifying and improving the functional characterizations of such genes was one of the objectives of the Gene Atlas experiment. Genes with single-copy orthologs in *A. thaliana* and consistent expression were significantly enriched in transcription factors (n = 501, 8.3%; Fisher’s exact test odds ratio = 1.507, *P* = 4.26e-13), suggesting that potential regulators of different biological processes are strongly conserved across the plant species (Keightley and Hill 1990). These observed evolutionarily conserved expression patterns inform functional details that complement direct sequence data comparisons.

In contrast to these ortholog-constrained analyses, co-expression analyses are agnostic to orthology, which dramatically increases the number of genes that can be analyzed, providing a broader perspective on gene expression regulatory evolution. For example, multidimensional scaling and hierarchical clustering revealed that phylogenetically neighboring species have more similar expression profiles across tissues and nitrogen treatments than more distantly related species (Mantel *R* > 0.63, *P* < 0.04) (**Fig. 2**). However, the phylogenetic signal of co-expression was dwarfed by variation among tissues, where far more of the total co-expression clustering across nitrogen source treatments was driven by patterns among tissues than genetic distance among species (tissues correlated with the first canonical correspondence analysis axis, which explains 41.46% of the variation), suggesting that genes in closely related species exhibit similar transcriptional profiles across tissues and conditions likely owing to the accumulation of evolutionarily conserved regulatory elements.

**Figure 2.**
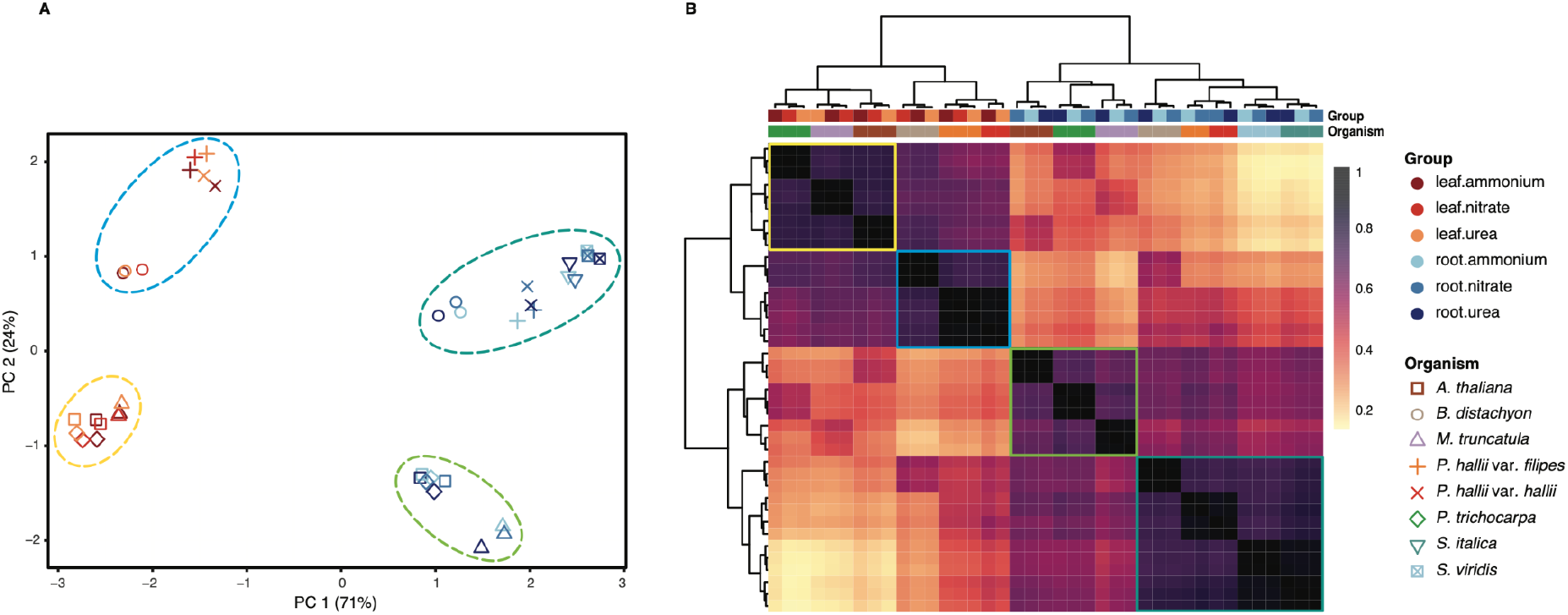
Global patterns of gene expression across eight vascular plants. Multidimensional scaling based on the expression of 2,066 single-copy orthologous genes in two tissues and three nitrogen treatment conditions show predominant clustering first by tissues and then by clade (mono-, dicots) (A). Hierarchical clustering based on Pearson correlation coefficients of log_2_ transformed normalized expression data (B).

### Patterns of tissue-specific gene expression across 18 species and >400M years of plant evolution

Tissue-specific expression complements global co-expression analyses by defining potential gene function associated with an organ or tissue. The major drawback of this approach results from morphological differences among species. For example, in *Chlamydomonas*, a single-celled organism, transcriptionally active genes in a given condition represent expressed genes in the organism as a whole, whereas multicellular organisms exhibit gene expression variation across different cell subtypes. Furthermore, the mosses sampled here lack root systems, flowers, seeds or easily sampled reproductive organs. Even the far more closely related flowering plants have functionally divergent homologous structures, such as root nodules, panicles, florets, sepals, and rhizomes. As such, analysis of tissue-specific expression must be somewhat phylogenetically constrained and condensed into large-scale functional tissue types (**Fig. 1**).

Our data suggest that large proportions of annotated genes (27-68%, *M* = 44.7; *SD* = 12.7) are commonly expressed (FPKM > 1) in multiple tissues (**Supplemental Data 6**), confirming that many genes serve multiple functions across tissues and environments. However, there was considerable among-tissue variation across species (ANOVA *F* = 70.01, *df* = 16, *P* < 2e-16) where gene expression is driven by variation among tissues or conditions (**Supplemental Data 6**). Such variably expressed genes may have evolved diverse functions depending on the regulatory environment across cell types.

Despite considerable across-tissue expression, we observed 220,218 (32.1%) of all genes with high expression specificity to a single tissue or condition. To identify genes exhibiting such strong tissue or condition specific expression, we used the Tau method (Yanai et al. 2005) which accounts for the number of unique sample types and produces consistently robust results with highest correlation between datasets of varying sizes (Kryuchkova-Mostacci and Robinson-Rechavi 2017). Using this method, we identified genes specific to (1) reproductive and root tissue in *S. italica*, (2) leaf, inflorescence, and whole floret in switchgrass, (3) leaf, leaf blade, dry seed, and imbibed seed in *S. bicolor*, and (4) stem and flower related gene sets in *Brachypodium*. Of all the standard plant tissues, stem and leaf had the fewest uniquely expressed genes (two-tailed unpaired Welch’s t-test, *P* = 8.338e-06) while roots followed by flower tissues were most unique (two-tailed unpaired Welch’s t-test, *P* = 2.547e-10). Groups of genes with greater expression proclivity towards spores, protonema and leaflet were recognized in *Physcomitrium*; drought and high temperature in *Sphagnum*; and towards seed, root tip, lateral root, and nodules in soybean (**Supplemental Data 7, 8**). These gene sets were largely overrepresented in GO biological processes known for each tissue or condition (**Supplemental Data 9**). Genes and their promoter regions with such marked expression specificity represent valuable tissue-specific reporters and targets for plant genetic engineering applications.

### Transcription modulation across developmental stages

Developmental time-courses represent a particularly powerful experiment to understand gene function and the dynamics of transcript abundance. As an example of such a time course, we evaluated the regulation of gene expression in leaf tissue in five developmental stages of *Sorghum bicolor* (juvenile, vegetative, floral initiation, anthesis and grain maturity). Overall, we identified 13,992 unique DEGs (*n* total annotated genes= 34,211) across the five developmental stages (**Fig. 3A, 3B, 3C**). KEGG pathway enrichments of up-regulated differentially expressed genes were largely consistent with physiological expectations: photosynthesis, carbohydrate and N metabolism terms were overrepresented in juvenile/vegetative stages (*P* < 0.05, hypergeometric test), floral initiation/anthesis stages were enriched in reproductive organ development and hormone signal transduction, and grain maturity stage were enriched for amino acid metabolism and transport, and zeatin and tyrosine metabolism (**Fig. 3D, 3E**). We observed the enrichment pattern to be reversed among downregulated genes in different stages, e.g., plant-pathogen interaction and plant hormone signal transduction were suppressed in juvenile and vegetative stages whereas photosynthesis, carbohydrate and N metabolism related pathways were among those suppressed in late developmental stages (**Supplemental Fig. 3**). These overrepresented pathways among DEGs at each stage illustrate the key biological events over the growing season, e.g., as juveniles the *S. bicolor* are collecting energy to increase the biomass, and as they flower and mature, they express defense mechanisms, and finally, with grain maturity, they reduce photosynthesis and slow down nutrient acquisition. The *S. bicolor* dataset provides an example of high-resolution characterization of gene expression changes and insight into the molecular responses of the plant across developmental stages represented by the Gene Atlas dataset.

**Figure 3.**
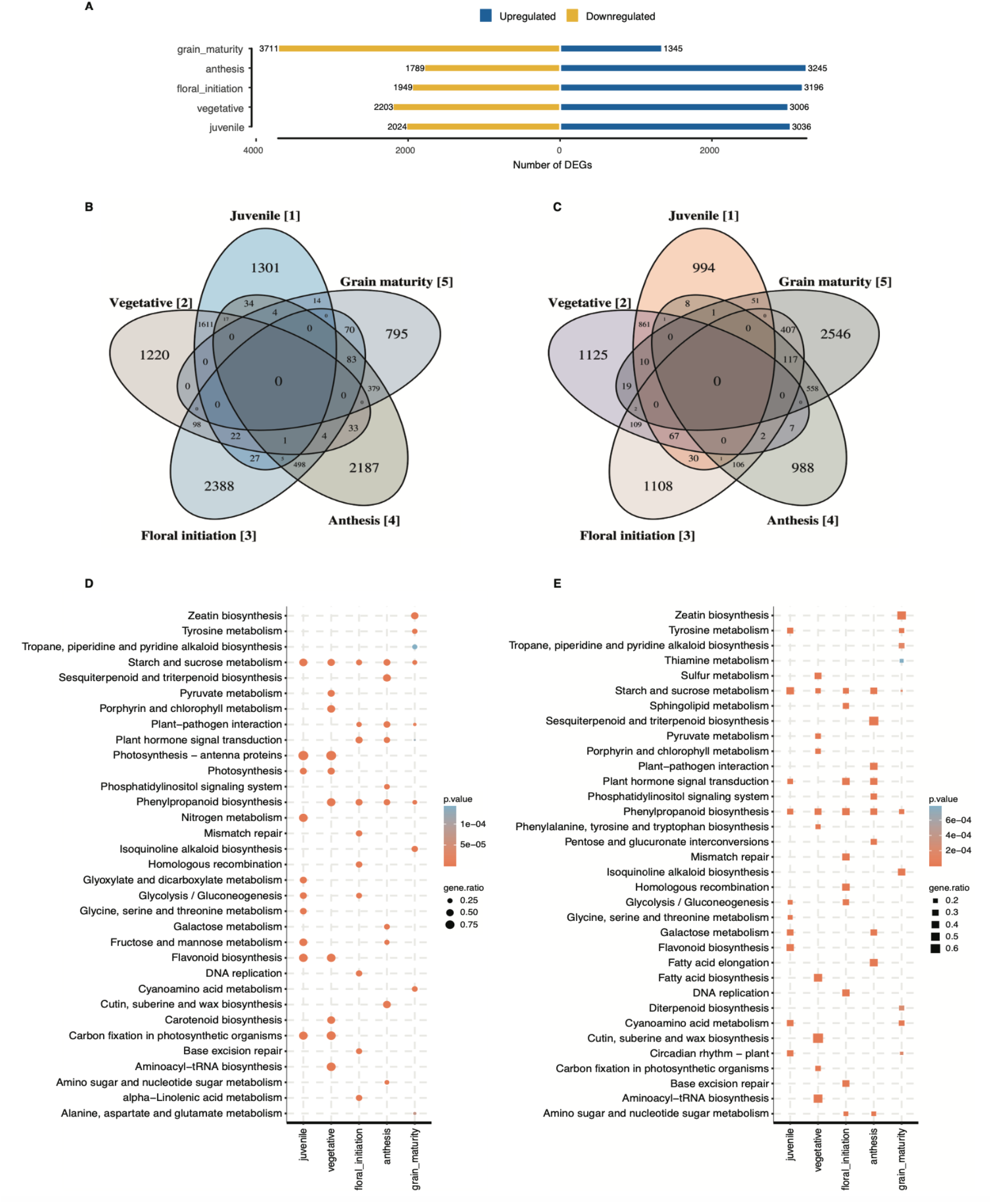
Differentially expressed gene comparison across five developmental stages in *Sorghum bicolor*. Numbers of differentially expressed genes across developmental stages (A). Venn diagrams of up-regulated (B) and down-regulated genes that are unique and shared between developmental stages (C). Top 10 KEGG metabolic pathway enrichments (*P* <.05, hypergeometric test) of up-regulated differentially expressed genes in each of the five developmental stages (D) and upregulated genes unique to each stage (E). ‘gene.ratio’ represents the ratio of number of DEGs over the number of genes annotated specific to the pathway.

### Transcriptional responses to different N sources

Tissue-specific gene expression regulatory responses to environmental cues are often evolutionarily conserved. These conserved responses offer a framework to test hypotheses about gene function as it relates to environmental sensitivity. A particularly powerful experiment adjusts the amount and type of necessary resource available. Drought, light and nutrient availability manipulations have provided strong evidence for gene function across the diversity of plants (Faye et al. 2022); (Zhang et al. 2021); (Huang, Zhao, and Chory 2019); (Swift et al. 2020); (Y. Li et al. 2022). In addition to providing evidence for the function of specific candidate genes’ responses to environmental stimuli, highly controlled manipulations, like our nitrogen source experiments, offer a framework to compare the relative roles of gene families and molecular pathways.

To understand gene expression underpinnings of N metabolism, we contrasted transcript abundance in aboveground and root tissues of each Gene Atlas species (where available, see **Fig. 1**) grown on N from three sources: urea, ammonium (NH_4_^+^), and nitrate (NO_3_^-^) (**Supplemental Data 10**). Since our experiments had similar statistical power and biological replicates among species and conditions, the total number of DEGs is a strong indicator of the transcriptional effects of different N sources. The most striking patterns were those related to tissue-specific gene expression variation within genotypes (**Fig. 4A, 4B**). For example, the root transcriptome was more responsive than aboveground tissues in all eudicot genotypes (Mann-Whitney U-test, *P* = 5e-04) except *Arabidopsis* (two-tailed unpaired Welch’s t-test, *P* = 0.4526). We observed consistent enrichments of N metabolism pathway genes among differentially expressed genes between treatments across many species, which demonstrates that this experiment elicits molecular responses of genes with homologs in genetic model species.

**Figure 4.**
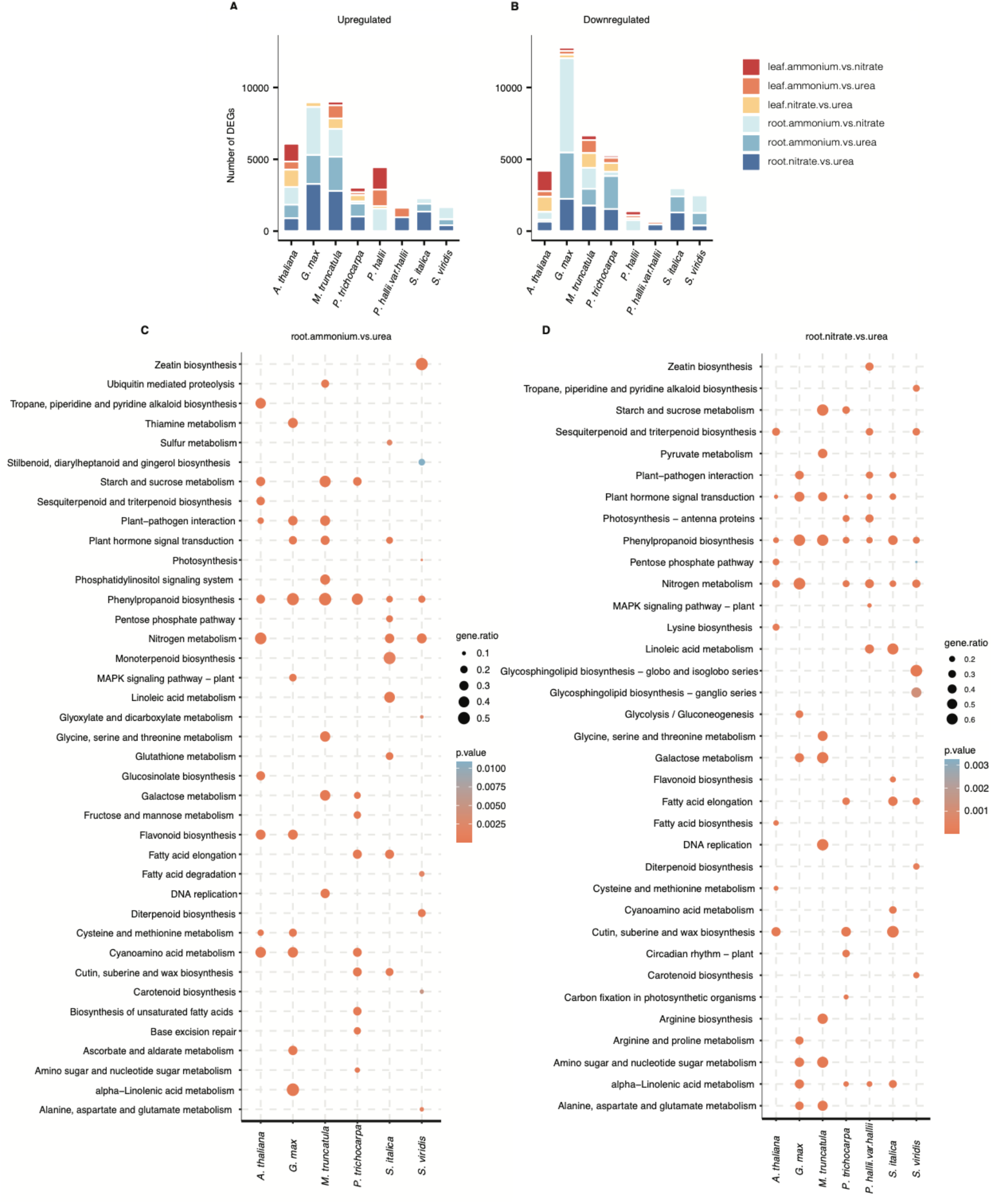
Transcriptional response of Gene Atlas plants towards NH_4_^+^ and NO_3_^-^ compared to urea as the sole nitrogen source in root and leaf tissues. Numbers of genes differentially upregulated (A) and numbers of genes differentially downregulated in response to changing nitrogen regime (B). Top 10 KEGG metabolic pathway enrichments (*P* <.05, hypergeometric test) in up-regulated differentially expressed genes in roots from Gene Atlas plants in ammonia vs. urea (C) and nitrate vs. urea treatment comparisons (D). ‘gene.ratio’ represents the ratio of number of DEGs over the number of genes annotated specific to the pathway.

Despite the power of discovering enriched groups of genes with similar and expected functional annotations, a major goal of the Gene Atlas is to provide a framework to discover novel gene functions and interactions. As such, we were excited to find starch and sucrose metabolism, and phenylpropanoid biosynthesis pathways overrepresented in upregulated DEGs. Indeed, many of the DEGs we identified in pairwise comparisons between N-sources are not directly involved in N metabolism. For example, genes associated with plant-pathogen interaction, plant hormone signal transduction, and carbohydrate metabolism were abundant (**Fig. 4C, 4D, Supplemental Fig. 4**). Similar observations were reported previously in *Sorghum* genotypes with varying N-stress tolerance subjected to N-limiting conditions (Gelli et al. 2014). Notably, nitrogen and amino acid metabolism-related pathways were overrepresented mainly in DEGs in nitrate vs. urea comparison. Such comparisons highlight differences in plant’s response to NO_3_^-^ as a sole N source compared to NH_4_^+^ at the metabolic level.

## INFERRING GENE FUNCTION FROM PATTERNS OF GENE EXPRESSION

### Variation in co-expression network topologies

Genes with similar expression patterns across diverse environmental conditions and tissues tend to serve similar biological functions across distantly related species and can be detected by co-expression clustering algorithms. For example, clusters of genes associated with a specific tissue or condition may be crucial for plant development or response to environmental cues. These strongly conserved tissue- and treatment-specific expression patterns facilitate biological gene function extrapolating from expression studies in one organism to close phylogenetic neighbors. To identify modules with such coherent expression patterns, we constructed 30 weighted gene co-expression networks (Langfelder and Horvath 2008) and 148 highly significant (min KME = 0.7, cut height = 0.25) co-expression modules within species and across different sets of tissues and conditions. Of these, 21 modules were significantly correlated with stress treatments (i.e., heat, cold, drought, salt, and wound stresses), 10 with N treatments, and 33 with other experimental conditions (**Supplemental Data 11**). Tissue-specific modules were also very common, e.g., root tissue-specific modules (n = 11), contained genes with GO terms enriched in responses to stimulus, oxidation-reduction process (Manzano et al. 2014; Passaia et al. 2014) and hydrogen peroxide metabolism (Ma et al. 2014) that are relevant to root functions (Bruex et al. 2012; Kogawara et al. 2014; W. Li, Lan, and 3.948 2015; Loreti et al. 2005). Leaf specific modules (n = 11) were enriched for phototropism, thylakoid membrane organization, pigment biosynthetic process, phototropism, and carbon fixation (**Supplemental Data 12**), suggesting that genes within the same module are associated with the same or interconnected biological functions.

### Inferring transcription factor functions from co-expressed genes

Genes showing highest connectivity with neighboring genes within a module, referred to as hub genes, are likely involved in preserving multi-gene regulatory variation and thus network integrity, potentially as trans-regulatory elements like transcription factors. We determined the top 10 most highly connected hub genes within each module. Across all the co-expression networks 87 hub genes belonged to transcription factor (TF) families (via PlantTFDB; (Jin et al. 2017)) (**Supplemental Data 13**), a slight but not significant enrichment of TFs relative to the genomic background (% hub TFs = 6.21%, background TFs = 5.23%, Fisher’s exact test odds ratio = 0.834, *P* = 0.104). TFs with many connections are presumed to be most influential in regulating the expression of modular genes in co-expression networks (Mukhtar et al. 2011). Under this premise, we further explored the overrepresented TF families among the hub genes. Most represented TF families in N treatment modules were MYB, WRKY, and NAC. Similar observations were made by Canales *et al*. (Canales et al. 2014) from Arabidopsis root transcriptomic data generated under contrasting N conditions. As shown in previous studies (Ghazalpour et al. 2006; Horvath et al. 2006; Liu, He, and Deng 2021; Miller, Oldham, and Geschwind 2008; Torkamani et al. 2010; Voineagu et al. 2011), hub genes play key roles in orchestrating module behavior and provide a specific focus for investigations into trait or condition related modules.

### Expression derived function descriptions (EDFD)

To evaluate how well the predicted gene function descriptions of Gene Atlas plants illustrate validated gene functions, we categorized currently assigned functional descriptions available at Phytozome as genes with good (GGF) and poor (GPF) function descriptions using an augmented dictionary lookup approach that incorporates weighting for negative, positive, and adversative keywords. Overall, 16% to 56% of the functional descriptions are GPF across the plants, with a large percentage of such genes not having a known function (**Supplemental Fig. 1**) (Gollery et al. 2006; Rhee and Mutwil 2014). We then assigned EDFD to the two subsets using results from tissue and condition specific expression groups, DEGs unique to a single contrast and co-expression network analysis along with ortholog function descriptions derived from nearest phylogenetic neighbors (see Methods).

Using this method, we added additional biological information to an average of 40.6% (*SD* = 12.6) of genes (excluding orthology based function descriptions) in these plant genomes (**Table 2; Supplemental Data 14**). For example, in the case of *S. bicolor*, 5,357 (15.65% of the total) genes lacked sequence homology-based function descriptions, 24,406 had good functional descriptions while overall 9,723 had poor descriptions. Gene Atlas expression-based functional descriptions were assigned to a total of 20,259 genes, of which 14,891 (43.63%of total annotated genes) had good functional descriptions and 5,368 (15.73%) had poor descriptions. To verify the reliability of the assigned functional associations, GO enrichment analysis of genes assigned with descriptions based on leaf and root samples was performed. We observed significant enrichment for photosynthesis, chloroplast organization, chlorophyll biosynthetic process and plastid translation (*P* <.05, Fisher’s exact test) in leaf related EDFDs; and cell wall loosening (Somssich, Khan, and Persson 2016), water transport and xyloglucan biosynthetic process (Peña et al. 2012) (*P* <.05, Fisher’s exact test) in root related EDFDs. Similar analysis in *Brachypodium* genes with assigned descriptions based on abiotic stress experiments (i.e., cold, heat, drought, and salt stress) showed significant enrichment for regulation of cellular response to alkaline pH, response to cold, heat, and positive regulation of response to oxidative stress (*P* <.05, Fisher’s exact test). Likewise, among genes annotated based on flower samples, specification of floral organ identity, fruit wall development and sporopollenin biosynthetic process were among the top enriched GO terms (*P* <.05, Fisher’s exact test). These results indicate that the assigned functional descriptions show strong biological role predictability and the approach here aids in expanding our current understanding of plant gene functions.

**Table 2.** Summary of assigned expression derived function descriptions (EDFD) to Gene Atlas. Number of annotated genes and the percentage of genes with good function descriptions (GGF), poor function descriptions (GPF) categorized using an augmented dictionary lookup approach that incorporates weighting for negative, positive, and adversative keywords and percentage of genes assigned with expression derived function descriptions.

| Genome | n.genes | % GGF | % assigned (GGF) | % GPF | % assigned (GPF) | n.assigned | % assigned (total) |
| --- | --- | --- | --- | --- | --- | --- | --- |
| *A. thaliana* | 27,416 | 83.86 | 32.43 | 16.14 | 5.573 | 10,418 | 38.00 |
| *B. distachyon* | 34,310 | 72.56 | 34.17 | 27.44 | 13.19 | 16,247 | 47.35 |
| *C. reinhardtii* | 17,741 | 43.08 | 12.6 | 56.92 | 15.98 | 5,070 | 28.58 |
| *E. grandis* | 36,349 | 79.74 | 28.63 | 20.26 | 3.997 | 11,858 | 32.62 |
| *G. max* | 52,872 | 80.37 | 47.25 | 19.63 | 11.33 | 30,971 | 58.58 |
| *K. fedtschenkoi* | 30,964 | 82.01 | 39.56 | 17.99 | 7.644 | 14,615 | 47.20 |
| *M. truncatula* | 50,894 | 67.94 | 23.66 | 32.06 | 5.285 | 14,731 | 28.94 |
| *P. hallii* | 33,805 | 72.65 | 34.97 | 27.35 | 8.656 | 14,746 | 43.62 |
| *P. hallii*HAL | 33,263 | 73.36 | 31.16 | 26.64 | 7.946 | 13,007 | 39.10 |
| *P. patens* | 32,926 | 55.44 | 19.35 | 44.56 | 14.06 | 11,003 | 33.42 |
| *P. trichocarpa* | 34,699 | 82.31 | 39.57 | 17.69 | 7.997 | 16,507 | 47.57 |
| *P. virgatum* | 80,278 | 69.2 | 39.73 | 30.8 | 14.15 | 43,251 | 53.88 |
| *S. bicolor* | 34,129 | 71.51 | 43.63 | 28.49 | 15.73 | 20,259 | 59.36 |
| *S. bicolor*Rio | 35,490 | 69.16 | 15.04 | 30.84 | 4.765 | 7,029 | 19.81 |
| *S. fallax* | 25,100 | 78.31 | 32.36 | 21.69 | 9.183 | 10,427 | 41.54 |
| *S. italica* | 34,584 | 77 | 39.37 | 23 | 10.78 | 17,344 | 50.15 |
| *S. viridis* | 38,334 | 70.43 | 35.99 | 29.57 | 12.74 | 18,680 | 48.73 |
| *L. albus* | 38,258 | 78.17 | 11.01 | 21.83 | 2.415 | 5,138 | 13.43 |

To help investigators target important genes for additional functional studies, we ranked the biological relevance of genes using a scoring methodology based on expression patterns of genes identified using tissue/condition specificity, differential expression, co-expression, hub status in a co-expression module and consensus expression across species. Gene orthologs with similar expression profiles in two or more species were given additional scores derived from the phylogenetic distance, i.e., larger the divergence time higher the score (see Methods). We identified a total of 656 top ranked genes across Gene Atlas plants (604 have orthologs in ≥5 plants; 40 of which have orthologs in ≥10 of evaluated plants) that have poor functional information but with the potential to improve our understanding of plant biology and form a list of prioritized targets for future experimental investigations (**Table 3; Supplemental Data 15**).

**Table 3.** Prioritized top ranked genes with poor functional descriptions for future experimental investigations. Genes were given scores based on expression patterns identified from i) unique differential expression in a single contrast, ii) tissue/condition specific expression and iii) biologically relevant co-expression modules (each given a score of 2) while hub genes in a co-expression module were given a score of 4. Gene orthologs with similar expression profiles were given additional scores derived from the phylogenetic distance. Total score was calculated as the aggregate of individual scores. Top ranked genes (two per species) are represented here.

| Organism | Gene ID | Score |  |  |  |  |  | Arabidopsis orthologs |
| --- | --- | --- | --- | --- | --- | --- | --- | --- |
|  |  | Differential expression | Condition specific expression | Co-expression | Hub gene | Consensus expression | Total |  |
| *A. thaliana* | AT2G20080 | 2 | 2 | 2 | 0 | 21.91 | 27.91 | AT2G20080 |
|  | AT4G28840 | 0 | 2 | 2 | 0 | 21.91 | 25.91 | AT4G28840 |
| *B. distachyon* | Bradi1g38210 | 0 | 2 | 2 | 0 | 10.7 | 14.7 | AT2G42760 |
|  | Bradi2g23445 | 0 | 0 | 2 | 0 | 11.69 | 13.69 | AT5G02090; AT2G37750 |
| *C. reinhardtii* | Cre02.g078550 | 0 | 2 | 2 | 4 | 0 | 8 |  |
|  | Cre02.g092700 | 0 | 2 | 2 | 4 | 0 | 8 |  |
| *E. grandis* | Eucgr.B00604 | 0 | 2 | 2 | 0 | 16.57 | 20.57 | AT5G08050 |
|  | Eucgr.F01122 | 0 | 2 | 2 | 0 | 17.12 | 21.12 |  |
| *G. max* | Glyma.13G227500 | 0 | 0 | 2 | 0 | 20.28 | 44.56 | AT1G33055 |
|  | Glyma.16G013600 | 0 | 0 | 2 | 0 | 20.84 | 45.68 | AT3G14280 |
| *K. fedtschenkoi* | Kaladp0065s0016 | 0 | 0 | 2 | 0 | 19.79 | 21.79 | AT4G28840; AT2G20080 |
|  | Kaladp0965s0006 | 2 | 0 | 2 | 0 | 26.23 | 30.23 | AT2G30230; AT1G06980 |
| *M. truncatula* | Medtr2g079300 | 2 | 2 | 2 | 0 | 18.11 | 24.11 |  |
|  | Medtr3g031140 | 0 | 2 | 2 | 0 | 27.28 | 31.28 | AT2G30230; AT1G06980 |
| *P. hallii* var. *filipes* | Pahal.3G090000 | 2 | 2 | 2 | 0 | 13.2 | 19.2 | AT5G02160 |
|  | Pahal.7G305700 | 2 | 2 | 2 | 0 | 12.05 | 18.05 | AT4G21445 |
| *P. hallii* var. *hallii* | PhHAL.3G160400 | 0 | 0 | 2 | 0 | 14.55 | 16.55 |  |
|  | PhHAL.5G229300 | 2 | 2 | 2 | 0 | 11.532 | 17.532 | AT5G62770; AT3G27880; AT1G23710; AT1G70420 |
| *P. patens* | Pp3c11_15500 | 2 | 2 | 2 | 4 | 0 | 10 |  |
|  | Pp3c13_2427 | 0 | 2 | 2 | 4 | 0 | 8 |  |
| *P. trichocarpa* | Potri.018G084100 | 2 | 2 | 2 | 0 | 19.79 | 25.79 | AT4G28840; AT2G20080 |
|  | Potri.003G193400 | 2 | 0 | 2 | 0 | 20.89 | 24.89 |  |
| *P. virgatum* | Pavir.5NG404000 | 0 | 0 | 2 | 0 | 13.64 | 15.64 |  |
|  | Pavir.2NG640501 | 0 | 2 | 2 | 0 | 9.72 | 13.72 | AT5G13720 |
| *S. bicolor* | Sobic.009G229000 | 0 | 0 | 2 | 4 | 13.35 | 19.35 | AT4G28840; AT2G20080 |
|  | Sobic.001G118400 | 2 | 2 | 2 | 0 | 10.44 | 16.44 | AT1G73885 |
| *S. bicolor*Rio | SbRio.08G154700 | 0 | 2 | 2 | 0 | 12.31 | 16.31 | AT5G08050 |
|  | SbRio.10G134000 | 0 | 2 | 2 | 0 | 12.05 | 16.05 | AT4G01150 |
| *S. angustifolium* | Sphfalx02G142200 | 2 | 2 | 2 | 4 | 0 | 10 |  |
|  | Sphfalx11G077900 | 2 | 2 | 0 | 0 | 4.5 | 8.5 | AT3G03341 |
| *S. italica* | Seita.9G407600 | 2 | 0 | 2 | 0 | 12.7053 | 16.7053 | AT1G63410; AT3G14260 |
|  | Seita.9G436900 | 0 | 2 | 2 | 0 | 12.5741 | 16.5741 | AT2G30230; AT1G06980 |
| *S. viridis* | Sevir.1G151100 | 2 | 0 | 2 | 0 | 13.2732 | 17.2732 | AT1G12320; AT1G62840; AT3G60780; AT2G45360 |
|  | Sevir.5G247600 | 0 | 0 | 2 | 0 | 13.2732 | 15.2732 | AT5G62770; AT3G27880; AT1G23710; AT1G70420 |

## DATA ACCESS

JGI Plant Gene Atlas data are currently hosted at two portals: i) JGI Plant Gene Atlas (https://plantgeneatlas.jgi.doe.gov), a dedicated portal provides bulk access to the data and user-specified queries of a single gene to multiple gene sets, lets users access differentially expressed genes, visualize gene ontology and pathway enrichments and track orthologs across the Gene Atlas plants; and ii) JGI’s plant portal, Phytozome (phytozome.jgi.doe.gov). Currently, Phytozome provides efficient tabular and graphical representation of co-expressed genes, pathway details, peptide, CDS and transcript sequence, protein homologs, plant family information and additionally genome browser view of gene models.

## CONCLUSIONS

Here we analyzed the transcriptional landscape of 18 plants from 2,090 RNA-seq datasets. To the best of our knowledge, it is the largest compendium of plant transcriptome data generated following standardized protocols across diverse plant species. These datasets enable JGI’s efforts to improve genome annotations especially related to conserved biological processes across the diversity of plants. Comparing orthologs among common gene sets between species allowed us to pinpoint and rank biologically relevant and evolutionarily conserved genes, demonstrating the potential of cross-species analysis from the transcriptome resource generated in this study. Furthermore, our results documented plant responses to varying N resources at the organ level and expression variation among developmental stages. These and other analyses highlight shared and varied gene expression regulatory evolution across plants.

The Gene Atlas datasets, along with the additional expression derived functional annotations, are valuable resources to the plant research community and provide targets, unknown or poorly described TFs, hub genes, and conserved genes, for functional studies that directly improve gene functional descriptions. We acknowledge that these functional associations are not definitive evidence of their functions, but we anticipate that they will be useful in directing future functional characterization experiments. We will continue to expand the Gene Atlas through standardized procedures to increase the specificity of these function descriptions. We strongly believe that results from this study and additional custom analyses on this resource will aid researchers in better understanding of roles of genes in their own experiments and get a better handle on biological processes at the system level.

## METHODS

### Plant growth and treatment conditions

#### *Glycine max* and *Medicago truncatula*

Plant seeds (*G. max* cv. Williams 82) were surface-sterilized, transferred to pots containing 3:1 vermiculite perlite. 2/3 seedlings were planted in each pot and grown until plants were 4 weeks in a growth chamber under 16 h-light/8 h-dark conditions, 26-23°C temperature maintained at 250 μmol m^−2^s^−1^. Plants for nitrogen experiment were watered with nutrient solution containing either 10 mM KNO_3_ (NO_3_^-^ plants) or 10 mM (NH_4_)3PO_4_ (NH_4_^+^ plants) or 10 mM urea (urea plants). We selected urea as a control condition for the counter ions, potassium, and phosphate, as the best compromise. The nutrient solutions were renewed every 3 days. After 4 weeks, different tissues (leaf, stem, root, shoot, shoot tip, root tip, lateral roots, etc) for N regimes and standard conditions were harvested. Plants under symbiotic conditions were watered with nutrient solution containing 0.5 mM NH_4_NO_3_ every other week. Subsequently, root nodules, roots, and trifoliate leaves under symbiotic conditions were collected and tissues from flower open and un-open were harvested from field grown plants.

#### Arabidopsis thaliana

Seeds were cold-stratified in water for 3 days and subsequently seeds were sown into 9 cm^2^ plastic pots (T.O. Plastics, Clearwater, FL, USA) filled with 2 parts Promix Biofungicide (Premier Tech, Riviére-du-Loup, QC, Canada) to 1 part Profile Field and Fairway (Profile, Buffalo Grove, IL, USA). Pots were placed in a growth chamber (22°C days/20°C nights, 14 h light at a photosynthetic photon flux density of 350 μmol m^−2^s^−1^), then thinned to 1 plant per pot containing Sunshine MVP potting mix (SunGro Horticulture) and transferred into a greenhouse at the University of Texas at Austin when rosettes had achieved 7-8 leaves. Plants supplemented with differing nitrogen source regimes (see *Glycine max*) were harvested after 30 days.

#### Brachypodium distachyon

Seeds (*B. distachyon* Bd21) were grown in Metro mix 360 soil in a growth chamber, under 12 h day and 12 h night conditions, maintained at 24°C/18°C, ~50% relative humidity; 150 μmol m^−2^s^−1^. Plants were watered once a day or every two days depending on the size of plants and soil conditions and fertilized twice a week (Tuesday and Friday) using Jack’s 15-16-17 at a concentration of 100 ppm. For the nitrogen source study, plants grown for 30 days under differing nitrogen source regimes (see *Glycine max*) were harvested.

For cold treatment experiment, Bd21 seeds were sown in soil without stratification. The germinated seeds were grown in a growth chamber under short day conditions (26°C 10 h light, 18°C 14 h dark) for 4 weeks and then moved to a cold room (4°C 10 h light, 4°C 14 h dark) for cold treatment. Whole shoots were harvested at different treatment time points and stored at −80°C for RNA extraction.

#### Chlamydomonas reinhardti

*C. reinhardtii* strain CC-1690 (also known as 21gr) was cultured at 24°C (agitated at 180 rpm at a photon flux density of 90 μmol m^−2^s^−1^ provided by cool white fluorescent bulbs at 4100 K and warm white fluorescent bulbs at 3000 K used in the ratio of 2:1) in tris acetate-phosphate (TAP) medium (Boyle et al. 2012). For growth in differing nitrogen sources, TAP medium was supplemented with (NH_4_)3PO_4_ or KNO_3_, or urea (see *Glycine max*). Cultures of strain CC-1690 were inoculated to 1 × 10^5^ cells ml^−1^ and collected for RNA at 1 × 10^6^ cells ml^−1^, when the growth rates of all cultures were identical. For assessing the impact of cell density, cultures were inoculated at 1 × 10^4^ cells ml^−1^ in replete medium and sampled at 5 × 10^5^ cells ml^−1^ and at each doubling thereafter until the culture reached a final density of 8 × 10^6^ cells ml^−1^.

#### Eucalyptus grandis

*E. grandis* samples were derived from tissues collected from clonal ramets of the genotype BRASUZ1 that was used to generate the *E. grandis* reference genome. Tissue samples were collected from three trees ca. 5 years old, and an adult tree ca. 8 years old at the time of sample collection, planted in experimental fields at Embrapa Genetic Resources and Biotechnology in Brasilia, Brazil (15.73 South, 47.90 West). RNA samples were prepared from adult leaves (completely developed), juvenile leaves (tender, thinner, not waxed), fruit buds, and developing cambium (from inside the tree bark). Plant material was collected from the field, immediately frozen in liquid nitrogen, and stored at −80°C until RNA extraction that followed an optimized CTAB-lithium chloride-based protocol (Inglis et al. 2018).

#### Kalanchoë fedtschenkoi

Four-week-old *K. fedtschenkoi* plants (accession ORNL M2) were grown under a 250 mmol m^−2^s^−1^ white light with a 12 h light (25°C)/12 h dark (18°C) cycle and were used as starting plant material for eight different experiments (i.e., circadian, metabolite, temperature, drought, light intensity, light quality, nitrogen utilization, and standard tissue). The experiments were conducted under day/night temperature regime of 25°C/18°C except the temperature experiment. For the circadian experiment, two sets of plants were grown under a 12 h light/12 h dark cycle and continuous lighting (250 mmol m^−2^s^−1^ white light), respectively, for seven days and then mature leaf samples (i.e., leaves 5-7 counting from the top of the plants) were collected every two hours over a 48 h period. For the metabolite experiment, plants were grown under an aerobic condition to prevent dark CO_2_ fixation and malate accumulation. This was accomplished by putting the plants in a sealed chamber with a closed air loop, through which air was continuously circulated. CO_2_ was subsequently continuously scrubbed from the air using a hydrated soda lime filter (LI-COR Biosciences, Lincoln NE) included in the loop. CO_2_ levels were monitored and maintained at an average of 3 ppm over the 12 h overnight aerobic treatment. Plants were removed from the aerobic condition just prior to the start of the daylight photoperiod. Mature leaves were harvested at 2 h intervals over the succeeding 24 h period (12 h light/12 h dark). For the temperature treatment, plants were grown under three different temperatures (8°C, 25°C and 37°C), respectively, for seven days. For drought treatments, plants were grown under three soil moisture conditions (40% ± 3% [control], 20% ± 3% [moderate drought] and 2% ± 3% [severe drought]), respectively, for 19 days. For the light intensity experiment, plants were grown under light intensity of 0 (darkness), 150 (low light) and 1000 (high light) mmol m^−2^s^−1^ for 48 h. For the light quality experiment, plants were grown under blue light (270 mmol m^−2^s^−1^), red light (280 mmol m^−2^s^−1^), far-red light (280 mmol m^−2^s^−1^) and constant darkness for 48 h. For the nitrogen utilization experiment, plants were treated with potassium sulfate (10 mM), ammonium sulfate (10 mM) and urea (5 mM), respectively, for four weeks. Immediately after the temperature, drought, light intensity, light quality and nitrogen utilization experiments, mature leaves were collected at two time points of dawn (2 h before the start of light period) and dusk (2 h before the start of dark period). For the nitrogen utilization experiment, root samples were also collected at dawn and dusk, respectively. For the standard tissue experiment, plants were grown in the greenhouse under a 12 h light/12 h dark cycle at Oak Ridge National Laboratory (Oak Ridge, TN) and five different tissue types (young leaf, young stem, mature stem, root, and flower) were collected at 10 am in the greenhouse.

#### Lupinus albus

RNA-seq data from cluster root samples were obtained from (Hufnagel et al. 2020).

#### Panicum virgatum

Vegetatively propagated Alamo AP13 plants were grown in pre-autoclaved MetroMix 300 substrate (Sungro^®^ Horticulture, http://www.sungro.com/) and grown in a walk-in growth chamber at 30/26°C day/night temperature with a 16 h photoperiod (250 μm^−2^ sec^−1^) for four months. Tissues were harvested at six developmental stages, including leaf development (VLD: V2), stem elongation (STE: E2 and E4), and reproductive phases (REP: R2, S2, and S6) (Moore et al. 1991).

For *P. virgatum* photoperiod experiment, four switchgrass genotypes, AP13, WBC, AP13, and VS16 plants were vegetatively propagated and grown in one-gallon pots with a 6:1:1 mixture of Promix:Turface:Profile soil at a growth chamber at the University of Texas at Austin. After one-week maintenance with a 30/25°C day/night temperature and 14L/10D photoperiod, plants from each genotype were divided into two groups and received LD (16L/8D) or SD (8L/16D) treatment in separate growth chambers. Fully emerged young leaves were simultaneously harvested from three individuals as three biological replicates after three-week LD and SD treatments. We collected two leaf tissues (2cm leaf tips and 2 cm leaf base) at two zeitgeber times (ZT1 and ZT17). All samples were immediately flash frozen in liquid nitrogen and stored at −80 °C for DNA and RNA extraction.

#### Panicum halli

The *P. hallii* FIL2 (var. *filipes*; Corpus Christi, TX; 27.65° N, 97.40° W) and *P. hallii* HAL2 (var. *hallii*; Austin, TX; 30.19° N, 97.87° W) were grown in 3.78 L pots at the University of TX Brackenridge Field Laboratory (Austin, Texas) in the greenhouse with mean daytime air temperature of 30°C and relative humidity of 65%. Plants supplemented with differing nitrogen source regimes (see *Glycine max*) were harvested after 30 days.

For *P. hallii* panicle samples, genotypes, HAL2 and FIL2, were grown in a growth chamber at University of Texas at Austin with 26°C day/22 °C night temperature and 12 h photoperiod. Plants were grown in 3.5 inches square pots with a 6:1:1 mixture of Promix:Turface:Profile soil. Young panicle tissues were collected under a dissection microscope and the developmental stages were determined according to the lengths (0.1-0.2 cm for D1 stage, 0.5-1 cm for D2 stage, 4.5-5.5 cm for D3 stage, and 9-11 cm for D4 stage). Tissues for D1 and D2 stages were taken from at least fifty plants and pooled for each biological replicate. Tissues for D3 and D4 stages were taken from at least fifteen plants and pooled for each biological replicate. All samples were harvested at 17:00-18:00 of the day and immediately flash frozen in liquid nitrogen. Three biological replicates for each stage were stored at −80°C for DNA and RNA extraction.

#### Physcomitrium patens

The protonemata cultures were systematically entrained by two successive weeks of culture prior to treatment to obtain a homogeneous culture as described in Perroud *et al*. (Perroud et al. 2018). In brief, BCD (Cove et al. 2009) or Knop medium (Reski and Abel 1985) were used to culture the moss. Solid medium (medium with 1% [w/v] agar) protonemal cultures were grown atop a cellophane film to allow tissue transfer for specific treatments (e.g., with hormones), and for ease of harvesting. Plates and flasks were cultivated at 22°C with a 16 h-light/8 h-dark regime under 60-80 μmol m^−2^s^−1^ white light (long-day conditions). All harvests were performed in the middle of the light photoperiod (+8 h of light in long day conditions) (Perroud et al. 2018; Fernandez-Pozo et al. 2020).

#### Populus trichocarpa

*Populus trichocarpa* (Nisqually-1) cuttings were potted in 4” X 4” X 5” containers containing 1:1 mix of peat and perlite. Plants were grown under 16 h-light/8 h-dark conditions, maintained at 20-23°C and an average of 235 μmol m^−2^s^−1^ to generate tissue for (1) standard tissues and (2) nitrogen source study. Plants for standard tissue experiment were watered with McCown’s woody plant nutrient solution and plants for nitrogen experiment were supplemented with either 10mM KNO_3_ (NO_3_^-^ plants) or 10mM (NH_4_)_3_PO_4_ (NH_4_^+^ plants) or 10 mM urea (urea plants). Once plants reached leaf plastochron index 15 (LPI-15), leaf, stem, root, and bud tissues were harvested and immediately flash frozen in liquid nitrogen and stored at −80°C until further processing was done.

The plant material for the seasonal time course study was obtained from 2-year-old branches and apical buds (understood as the top bud of each branch) of 5-year-old hybrid poplar (*Populus tremula × alba* INRA 717 1B4) trees planted at the Centre for Plant Biotechnology and Genomics (CBGP) in Pozuelo de Alarcón, Madrid (3°49’W, 40°24’N), growing under natural conditions. Stem samples were collected weekly from November 7, 2014, to April 9, 2015. Buds were collected weekly from 13 January to 14 April 2015. For each time point, stem portions from 8 trees and 25 apical buds from 8 trees were pooled. RNA extraction was performed using the protocol described in (Ibañez et al. 2008). For the gene expression analysis, the weekly data were divided into groups named; fall, winter, and spring. Letter suffixes - “a, b, c, d, e” were added to group names representing “early,” “mid,”, “late”, “fortnight-1” or “fortnight-2” based on sampling dates within each season, following the Northern Meteorological Seasons dates.

#### *Setaria italica* and *Setaria viridis*

Seeds (*S. italica* B100 and *S. viridis* A10.1) were sown in flats (4×9 inserts/flat) containing Metro mix 360 soil and grown in a growth chamber, under 12 h day and 12 h night conditions, maintained at 31 °C/22°C, 50%-60% humidity; 450 μmol m^−2^s^−1^. Plants were watered once a day or every two days depending on the size of plants and soil conditions and fertilized twice a week (Tuesday and Friday) using Jack’s 15-16-17 at a concentration of 100 ppm. For light treatment experiments, plants were grown under continuous monochromatic light, blue: 6 μmol m^−2^s^−1,^ red: 50 μmol m^−2^s^−1,^ far-red: 80 μmol m^−2^s^−1^, respectively and watered with RO water every 3 days. Total aerial tissues were collected (at 9.30 AM) from 8-day old seedlings.

#### Sorghum bicolor

The reference line BTx623 was grown under 14 h day greenhouse conditions in topsoil to generate tissue for two separate experiments: (1) a nitrogen source study and (2) a tissue by developmental stage timecourse. For the nitrogen source study, plants grown under differing nitrogen source regimes (see *Glycine max*) were harvested at 30 days after emergence (DAE). For the tissue by developmental stage timecourse, plants were harvested at the juvenile stage (8 DAE), the vegetative stage (24 DAE), at floral initiation (44 DAE), at anthesis (65 DAE), and at grain maturity (96 DAE) and leaf, root, stem and reproductive structures as described in McCormick et al (Mccormick et al. 2017).

#### *Sorghum bicolor* var Rio

Genetic material for *S. bicolor* var Rio was obtained from a single seed source provided by W. Rooney at Texas A&M University. Plants were grown in greenhouse conditions and material for RNA extraction was collected at 6 biological stages: vegetative (5-leaf), floral initiation, flag leaf, anthesis, soft dough, and hard dough. Stages were identified based on biological characteristics defined in (Vanderlip and Reeves 1972). At every stage, whole plants were harvested, and the topmost fully developed leaf and topmost internode were collected. During the first 3 stages, meristems were isolated from the topmost internode while floral and seed tissues were collected after plants had flowered. All tissues were immediately placed in RNA Later and stored at 4°C prior to RNA extraction. See also (Cooper et al. 2019).

#### *Sphagnum angustifolium* (formally *S. fallax*)

*S. angustifolium* were grown on BCD agar media pH 6.5, ambient temperature (20°C) and 350 μmol m^−2^s^−1^ of photosynthetically active radiation (PAR) at a 12 h light/dark cycle for 2 months prior to initiation of experimental conditions. At 8 am on the morning of the treatments, *Sphagnum* plantlets were transferred to petri dishes with 15 ml of appropriate BCD liquid media and placed in a temperature-controlled growth cabinet. Excluding the dark treatment, all samples were kept under 350 PAR for the duration of the experiment. Morning treatment samples were harvested at noon. After each experiment the material was blotted dry, placed in a 15 mL Eppendorf tube, flash frozen in liquid nitrogen, and stored at −80°C until RNA extractions were completed.

For the control treatment, *Sphagnum* plants were placed in a 22.05 cm^2^ petri dish containing BCD media 6.5 pH and incubated in a growth cabinet at 20°C and ambient light 350 PAR. To test low pH gene expression, the sample was placed in a 22.05 cm^2^ petri dish containing 6.5 pH BCD media at 8 AM. Each hour, the pH was gradually decreased until the sample was transferred to 3.5 pH media at 11 AM. The samples were harvested at 12 PM. This treatment was repeated for the high pH experiment except the sample was gradually brought from 6.5 to 9.0 pH. Temperature experiments were controlled in growth cabinets plantlets in 22.05 cm^2^ petri dishes containing 6.5 pH BCD media. The high temperature treatment began at 20°C and over three hours, temperature was gradually increased to 40°C. The low temperature treatment began at 20°C and over three hours, was gradually decreased to 6°C. To test water loss effects on gene expression, plantlets were placed on dry plates (no BCD media) for the duration of the experiment. Dark effect on gene expression was tested by placing plant material in a BCD filled petri dish in complete darkness from 8 AM to 12 PM. To evaluate gene expression that is present during immature growth stages, a sporophyte was collected from the mother of the *S. angustifolium* pedigree and germinated on solid Knop medium under axenic tissue culture conditions. After 10 days of growth, plantlets were predominantly within the thalloid protonemata with rhizoid stage and flash frozen in LN2 until RNA extraction using CTAB lysis buffer and Spectrum Total Plant RNA kit.

### RNA extraction and sequencing

All tissues were immediately flash frozen in liquid nitrogen and stored at −80°C until further processing was done. Every harvest involved at least three independent biological replicates for each condition. High quality RNA was extracted mainly using standard Trizol-reagent based extraction (Z. Li and Trick 2005), exceptions noted above under individual species. The integrity and concentration of the RNA preparations were checked initially using Nano-Drop ND-1000 (Nano-Drop Technologies) and then by BioAnalyzer (Agilent Technologies). Plate-based RNA sample prep was performed on the PerkinElmer Sciclone NGS robotic liquid handling system using Illumina’s TruSeq Stranded mRNA HT sample prep kit utilizing poly-A selection of mRNA following the protocol outlined by Illumina in their user guide: http://support.illumina.com/sequencing/sequencing_kits/truseq_stranded_mrna_ht_sample_prep_kit.html, and with the following conditions: total RNA starting material was 1 μg per sample and 8 cycles of PCR was used for library amplification. The prepared libraries were then quantified by qPCR using the Kapa SYBR Fast Illumina Library Quantification Kit (Kapa Biosystems) and run on a Roche LightCycler 480 real-time PCR instrument. The quantified libraries were then prepared for sequencing on the Illumina HiSeq sequencing platform utilizing a TruSeq paired-end cluster kit, v4, and Illumina’s cBot instrument to generate a clustered flow cell for sequencing. Sequencing of the flow cell was performed on the Illumina HiSeq2500 sequencer using HiSeq TruSeq SBS sequencing kits, v4, following a 2×150 indexed run recipe. The same standardized protocols were used to prevent introduction of any batch effects among samples throughout the project.

### RNA-seq data normalization and differential gene expression analysis

Illumina RNA-seq 150 bp paired-end strand-specific reads were processed using custom Python scripts to trim adapter sequences and low-quality bases to obtain high quality (Q≥25) sequence data. Reads shorter than 50 bp after trimming were discarded. The processed high-quality RNA-seq reads were aligned to current reference genomes of Gene Atlas using GSNAP, a short read alignment program (Wu and Nacu 2010). HTSeq v1.99.2, a Python package was used to count reads mapped to annotated genes in the reference genome (Anders, Pyl, and Huber 2015).

Multiple steps for vetting libraries and identifying outliers were employed, including visualizing the multidimensional scaling plots to identify batch effects, if any, and outliers among the biological replicates were further identified based on Euclidean distance to the cluster center and the Pearson correlation coefficient, *r* ≥ 0.85. Libraries retained after QC and outlier-filtering steps were only considered for further analysis. Detected batch effects, if any, were accounted for using RUVSeq (v1.4.0) (Risso et al. 2014) with the residual RUVr approach. Fragments per kilobase of exon per million fragments mapped (FPKM) and transcripts per million (TPM) values were calculated for each gene by normalizing the read count data to both the length of the gene and the total number of mapped reads in the sample and considered as the metric for estimating gene expression levels (B. Li and Dewey 2011; Trapnell et al. 2011). Genes with low expression were filtered out, by requiring ≥2 relative log expression normalized counts in at least two samples for each gene. Differential gene expression analysis was performed using the DESeq2 package (v1.30.1) (Love, Huber, and Anders 2014) with adjusted *P*-value < 0.05 using the Benjamini & Hochberg method and an log_2_ fold change >1 as the statistical cutoff for differentially expressed genes.

### Co-expression network construction

Weighted gene co-expression networks were constructed using the WGCNA R package (v1.70.3) (Langfelder and Horvath 2008) with normalized expression data retained after filtering genes showing low expression levels (log_2_ values of expression <2). Subsets of samples belonging to specific experiments such as N study, developmental stages, or stress treatment, were used to construct multiple networks for each species. Subsetting samples reduces the noise and increases the functional connectivity and specificity within modules. We followed standard WGCNA network construction procedures for this analysis. Briefly, pairwise Pearson correlations between each gene pair was weighted by raising them to power (β). To select proper soft-thresholding power, the network topology for a range of powers was evaluated and appropriate power was chosen that ensured an approximate scale-free topology of the resulting network. The pairwise weighted matrix was transformed into a topological overlap measure (TOM). And the TOM-based dissimilarity measure (1 – TOM) was used for hierarchical clustering and initial module assignments were determined by using a dynamic tree-cutting algorithm. Pearson correlations between each gene and each module eigengene, referred to as a gene’s module membership, were calculated and module eigengene distance threshold of 0.25 was used to merge highly similar modules. These co-expression modules were assessed to determine their association with expression patterns distinct to a tissue or condition. Module eigengenes were associated with tissues or treatment conditions or developmental stages to gain insight into the role each module might play. These modules were visualized using igraph R package (v.1.2.6) (Gabor Csardi and Tamas Nepusz 2006) and in order to focus on relevant gene pair relationships, network depictions were limited to top 500 within-module gene-gene interactions as measured by topological overlap.

### GO and KEGG pathway enrichment analysis

GO enrichment analysis of DEGs, co-expression modules and genes in tissue and condition specific clusters was performed using topGO (v.2.42.0) (Alexa A and Rahnenfuhrer J 2016), an R Bioconductor package, to determine overrepresented GO categories across biological process (BP), cellular component (CC) and molecular function (MF) domains. Enrichment of GO terms was tested using Fisher’s exact test with *P* <0.05 considered as significant. KEGG (Kanehisa and Goto 2000) pathway enrichment analysis was also performed on those gene sets based on hypergeometric distribution tests and pathways with *P* <0.05 were considered as enriched.

### Categorization of function descriptions

An augmented dictionary lookup approach that incorporates weighting for positive (amplifiers), negative (including attenuators), and adversative keywords was adapted from sentiment analysis methodology to categorize gene function descriptions. We generated a custom dictionary from gene function descriptions of all Gene Atlas plants and used a modified valence shifters data table with sentimentr (v.2.9.0) (https://cran.r-project.org/web/packages/sentimentr), to obtain sentiment score. We empirically determined the minimum cutoff for sentiment score to classify gene descriptions as good (score > 0.3) and poor (score < 0.3) function descriptors.

### Identification of orthologous genes

OrthoFinder (v2.5.4) was used to identify orthologous genes across 18 Gene Atlas species using default parameters (Emms and Kelly 2019). OrthoFinder results were parsed to generate tables of orthologs for each species and genes with one-to-one ortholog relationships between species identified using rooted gene trees were further subsetted.

### Gene ranking method

To rank and prioritize genes by their biological relevance, genes with distinct expression patterns identified based on i) tissue/condition specificity, ii) unique DE in a single contrast were given a score of 2 for each method i.e., a gene was assigned a score of 4 if it were identified by two methods. These scores were augmented with co-expression network analysis (described above). Genes in biologically relevant modules were ranked (score=2) while hub genes in a co-expression module were ranked the highest (score=4). Also, gene orthologs with consensus expression pattern in two or more plants were given additional scores based on the phylogenetic distance between species (Zeng et al. 2014; Kumar et al. 2017) i.e., larger the divergence time higher the score (million years ago/100) (**Supplemental Data 16**). Final ranking of the genes was calculated as the aggregate of individual scores.

### System design and implementation

All statistical analyses and visualizations were performed using the R 4.0.3 Statistical Software (R Development Core Team 2011) and its web interface was developed using shiny (v1.7.1). Currently, Gene Atlas is deployed on a CentOS Linux server by employing Docker (version 19.03.11), an open platform for developing and running applications.

## Data availability

The RNA-seq data that support the findings of this study are available from the NCBI Sequence Read Archive (SRA) under accessions provided in **Supplemental Data 1**. To enable exploration of the transcriptome datasets for JGI Plant Gene Atlas v2.0, the data are hosted on Gene Atlas portal (https://plantgeneatlas.jgi.doe.gov) and JGI’s Phytozome plant portal. Documentation for data processing and downloadable data are available in the ‘Methods’ section (https://plantgeneatlas.jgi.doe.gov).

## Competing financial interests

The authors declare no competing financial interests.

**Correspondence and requests for materials** should be addressed to A.S. or J.S.

## ACKNOWLEDGMENTS

The work (proposal: 10.46936/10.25585/60000843) conducted by the U.S. Department of Energy Joint Genome Institute (https://ror.org/04xm1d337), a DOE Office of Science User Facility, is supported by the Office of Science of the U.S. Department of Energy operated under Contract No. DE-AC02-05CH11231.

The *Populus* work was partially supported by the U.S. Department of Energy under Contract to Oak Ridge National Laboratory. Oak Ridge National Laboratory is managed by UT-Battelle, LLC for the US Department of Energy under contract number DE-DE-AC05-00OR22725. Specific funding for the soybean transcriptome atlas was provided by a grant from the United Soybean Board (to GS). The switchgrass work was carried out under the support of the BioEnergy Science Center (BESC, a U.S. Department of Energy Bioenergy Research Center supported by the Office of Biological and Environmental Research in the DOE Office of Science, U.S. Department of Energy) and funded by the Samuel Roberts Noble Foundation. BH work was funded in part by U.S. DOE the Office of Biological and Environmental Research under grant number DE-SC0012629. The *Chlamydomonas* work is supported by the US Department of Energy Grant DE-FC02-02ER63421 and by the National Institutes of Health (NIH) R24 GM092473 to SM. The Eucalyptus work was supported by the Brazilian Federal District Research Foundation (FAP-DF) NEXTREE grant. The *Panicum hallii* work was supported by the DOE Office of Science, Office of Biological and Environmental Research (BER), grant no. DE-SC0008451 and DE-SC0021126 to TEJ. The sorghum work by JM laboratory was funded in part by the DOE Great Lakes Bioenergy Research Center (DOE BER grant DE-SC0018409). The *Setaria* work was funded by the DOE Office of Science under grant numbers DE-SC0018277 and DE-SC0008769. The *Kalanchoë* work was partially supported by the DOE Office of Science, Genomic Science Program under Award No. DE-SC0008834. Research related to Sphagnum was funded by DOE BER Early Career Research Program at Oak Ridge National Laboratory is managed by UT-Battelle, LLC, for the US DOE under contract number DE-AC05–00OR22725. Thanks to Daniel S. Rokhsar for involvement in the early formulation of Gene Atlas and helpful discussions.

## AUTHOR CONTRIBUTIONS

A.S. and C.P. conducted transcriptome data analyses. J.T.L. conducted metadata analysis. MSH carried out soybean and *Medicago* experiments. J.G., K.B., M.A., M.K., M.W., A.L., Jenifer J, L.S., K.A., M.Z., C.D. performed RNA-seq library preparation and sequencing. J.K. and S.D.G. conducted *Chlamydomonas* experiments. J.C., J.P., and D.G. maintains the data repository at Phytozome. J.W.J. and C.P. assembled genomes. S.S. conducted genome annotation at JGI. I.T-J., and M.U. conducted *Panicum virgatum* experiments. S.S.J. conducted *Populus* experiments and *Sphagnum* RNA extractions. D.C. and M.P. conducted *Populus* seasonal time course experiments. H.J., C.S., P.H., J.S., C.L., A.M. and S.C. conducted *Setaria* experiments and sample preparations. L.L. conducted Brachypodium cold treatment experiments. A.A.C. conducted *Sphagnum* experiments. B.W. conducted *Sorghum* N-treatment experiments. R.H. conducted *Kalanchoe* experiments. M.R.P. conducted *Eucalyptus* experiments. R.T. and K.S. conducted validation experiments for N-treatment. E.V.S. and X.W. conducted Arabidopsis, *P. hallii* and *P. virgatum* photoperiod experiments and sample preparations. A.M. conducted *P. virgatum* drought stress experiments and sample preparation. P.F.P. and F.B.H. analyzed *Physcomitrium* data. P.F.P., M.H. and S.A.R. provided *Physcomitrium* samples. D.S.R., D.G., D.T., D.W., E.A.C., E.K., G.S., G.A.T., I.B., J.S., J.M., J.V., S.A.R., S.S.M., T.B., T.E.J., T.C.M., X.Y., and Y.T. are principal investigators (alphabetical order). All authors read and approved the final manuscript.

A.S., J.T.L., and J.S. prepared the manuscript with input from all authors.

## SUPPLEMENTAL MATERIAL

**Supplemental Data 1**. Correlation among biological replicates in Gene Atlas species.

**Supplemental Data 2**. RNA-seq samples generated/analyzed in this study.

**Supplemental Data 3**. One-to-one orthologous genes across eight vascular plants.

**Supplemental Data 4**. One-to-one orthologous genes with consistent expression among species, but weak functional descriptions.

**Supplemental Data 5**.*Arabidopsis thaliana* (TAIR10) orthologs of genes with conserved expression patterns across Gene Atlas plants.

**Supplemental Data 6**. Percentage of genes commonly expressed in multiple tissues in Gene Atlas plants.

**Supplemental Data 7**. Genes with strong expression proclivity towards selected tissues/conditions in *G. max, P. patens* and *S. angustifolium*.

**Supplemental Data 8**. Genes with strong tissue/condition specific expression across Gene Atlas plants.

**Supplemental Data 9**. Overrepresented biological processes among genes with strong tissue/condition specific expression.

**Supplemental Data 10**. Differentially expressed genes in nitrogen treatment study.

**Supplemental Data 11**. Co-expression network module genes generated across different sets of tissues and conditions within Gene Atlas plants.

**Supplemental Data 12**. Overrepresented biological processes among co-expression network module gene sets generated across different sets of tissues and conditions within Gene Atlas plants.

**Supplemental Data 13**. List of hub genes in co-expression network modules.

**Supplemental Data 14**. Expression derived function descriptions (EDFD) assigned to annotated genes across Gene Atlas plants.

**Supplemental Data 15**. Prioritized top ranked genes with poor functional descriptions for future experimental investigations.

**Supplemental Data 16**. Estimates of divergence time between Gene Atlas species.

## SUPPLEMENTAL FIGURES

**Supplemental Figure 1.**
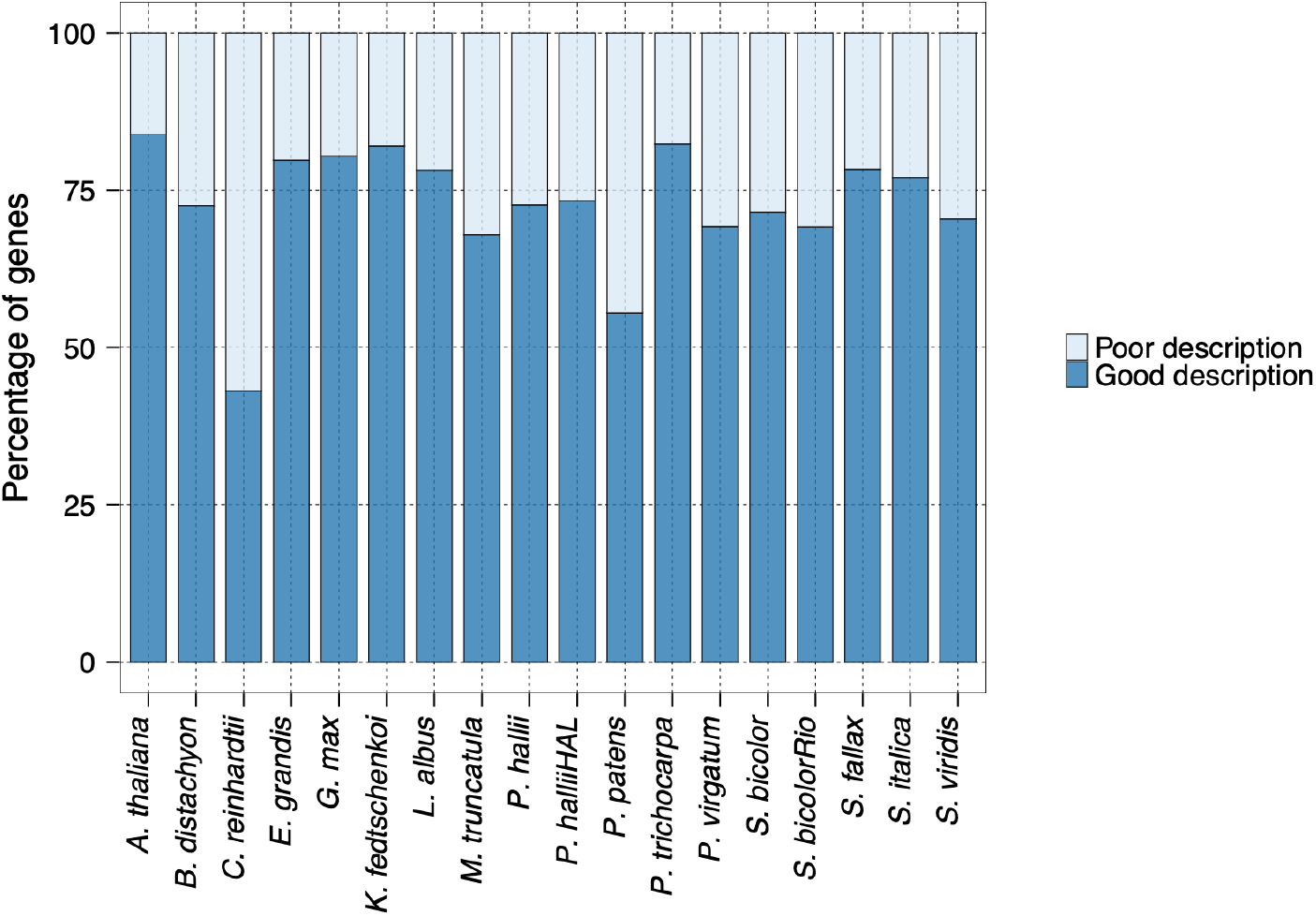
Classification of gene function descriptions. Percentage of genes with poor and good function descriptions categorized using an augmented dictionary lookup approach that incorporates weighting for negative, positive and adversative keywords.

**Supplemental Figure 2.**
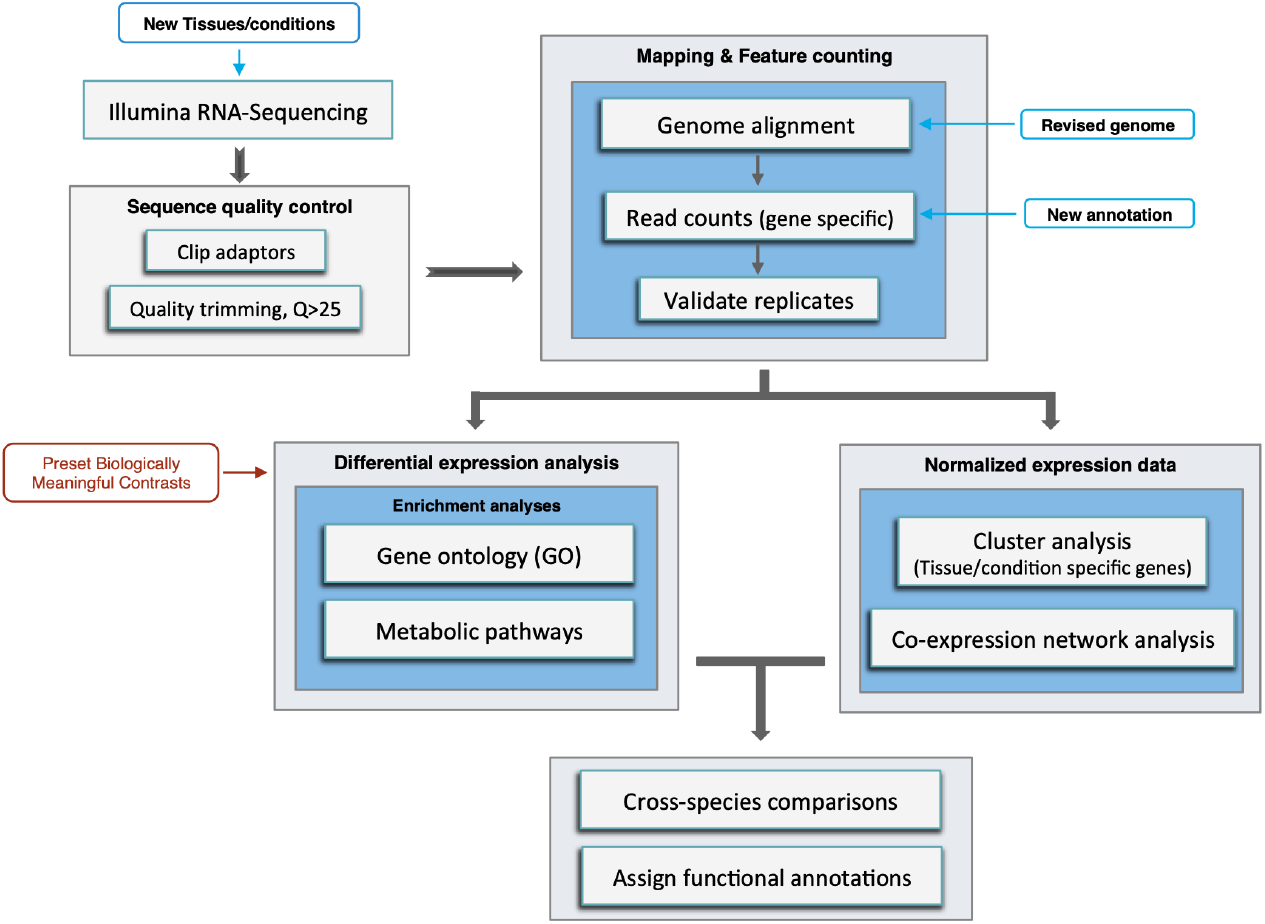
Plant Gene Atlas analysis flowchart. Pipeline representing methodology used to analyze RNA-seq data and assign experimentally derived functional annotations.

**Supplemental Figure 3.**
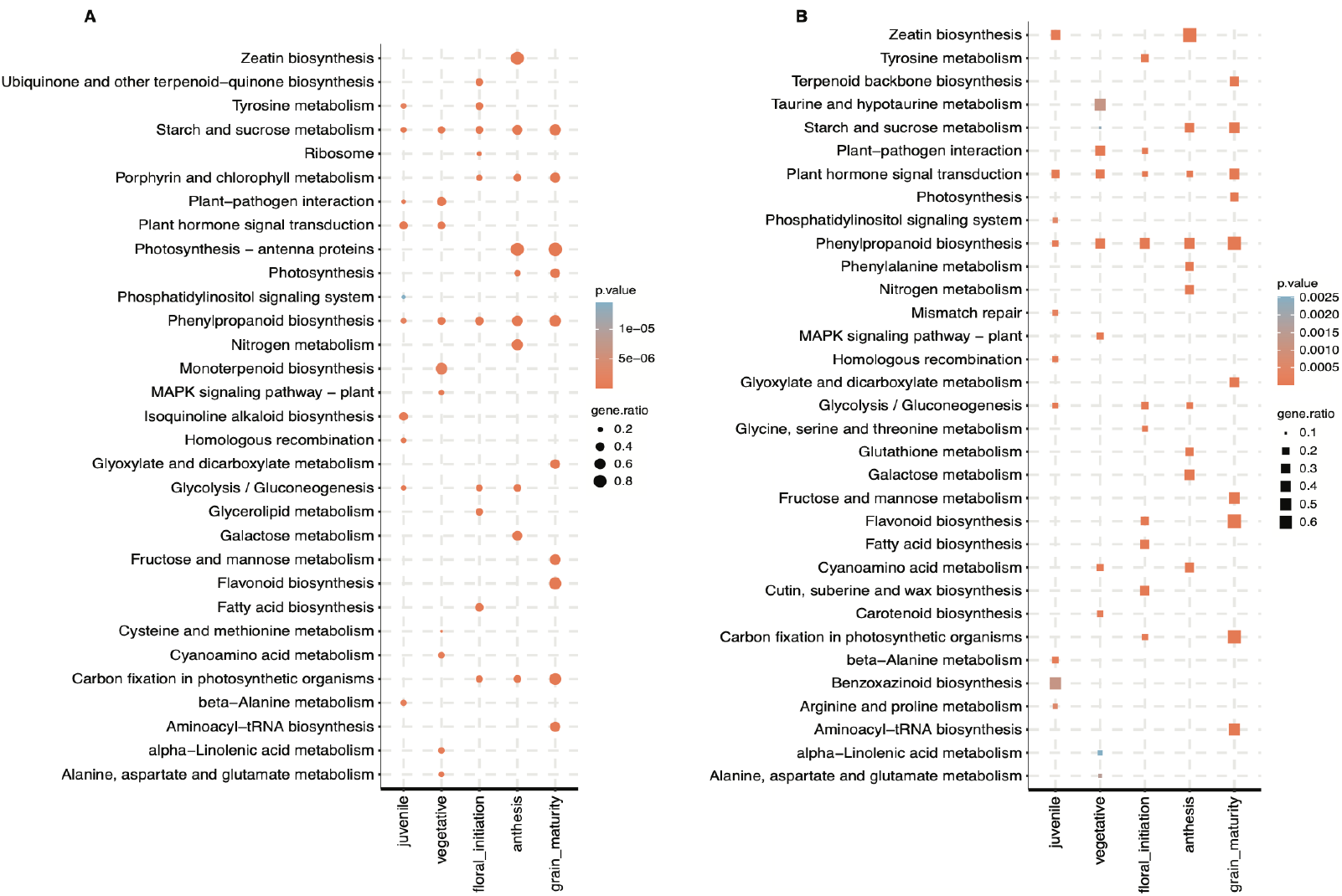
Differentially expressed gene comparison across five developmental stages in *Sorghum bicolor*. Top 10 KEGG metabolic pathway enrichments (*P* <.05, hypergeometric test) of down-regulated differentially expressed genes in each of the five developmental stages (A) and down-regulated genes unique to each stage (B). ‘gene.ratio’ represents the ratio of number of DEGs over the number of genes annotated specific to the pathway.

**Supplemental Figure 4.**
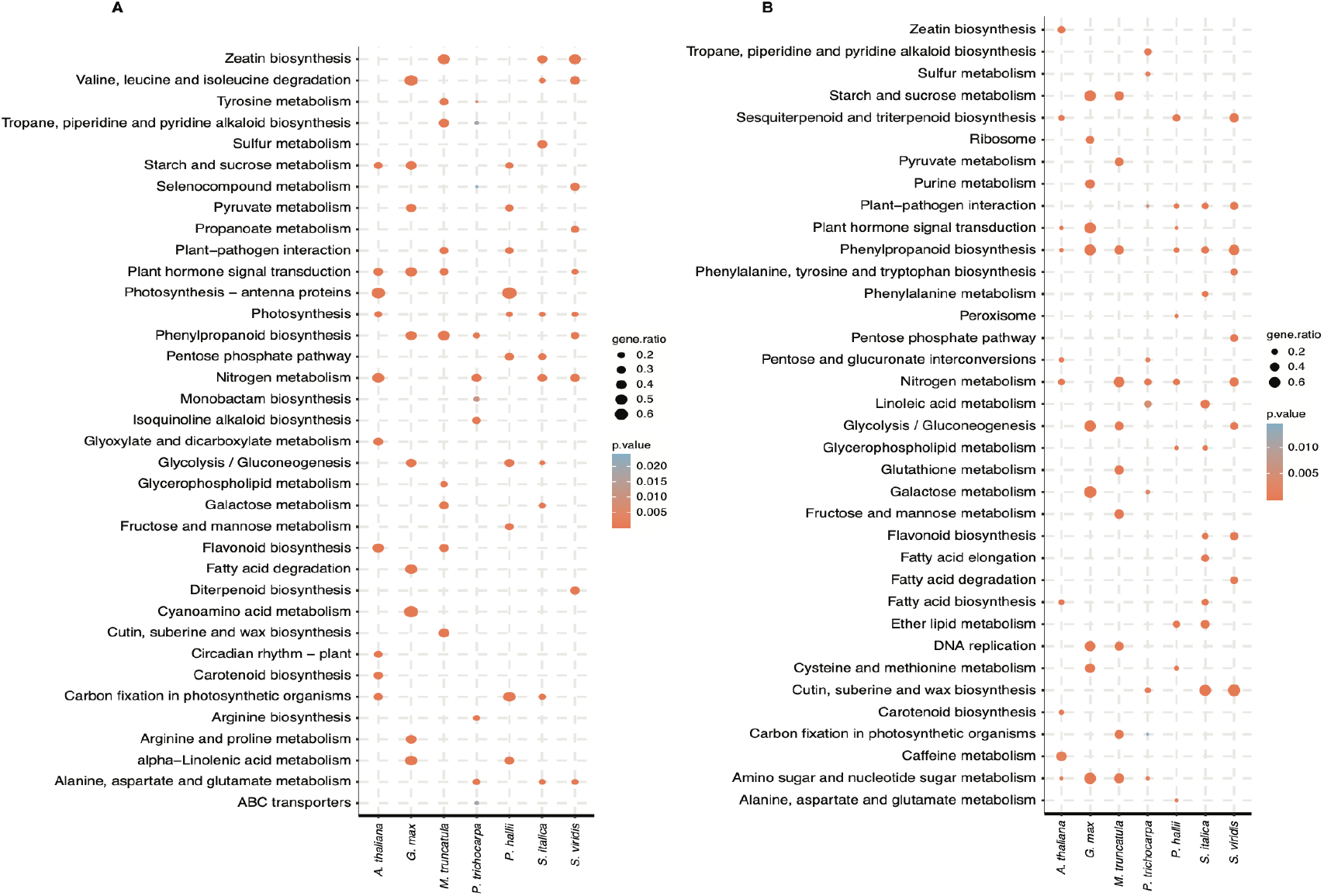
Transcriptional response of Gene Atlas plants towards NH_4_^+^ and NO_3_^-^ as the sole nitrogen source in root tissues. Top 10 KEGG metabolic pathway enrichments (*P* <.05, hypergeometric test) in up-regulated (A) and down-regulated (B) differentially expressed genes in roots from Gene Atlas plants in ammonia vs. nitrate comparison. ‘gene.ratio’ represents the ratio of number of DEGs over the number of genes annotated specific to the pathway.

